# PFOA exposure amplifies normal developmental gene expression programs in the African Killifish, *Nothobranchius furzeri*

**DOI:** 10.64898/2026.06.15.732418

**Authors:** Zainab Afzal, Charles Hatcher, Vandana Veershetty, Evan E Pittman, Deepak Kumar

## Abstract

Per- and polyfluoroalkyl substances (PFAS) are persistent environmental contaminants associated with developmental abnormalities and adverse health outcomes, yet it remains unclear whether PFAS exposure imposes novel transcriptional programs during development or perturbs endogenous developmental processes. Here, we continuously exposed African killifish (*Nothobranchius furzeri*) to an environmentally relevant concentration of perfluorooctanoic acid (PFOA) from egg laying through juvenile development to mimic prenatal-to-adolescent exposure and performed whole-transcriptome sequencing at two developmental stages. Despite four weeks of embryonic exposure, newly hatched juveniles, approximately equivalent to human infants, exhibited remarkably limited transcriptional responses, with only a few differentially expressed genes identified. In contrast, older juveniles, equivalent to human adolescents, exposed for eight weeks displayed a dramatic expansion of transcriptional perturbation, with approximately 30-fold more differentially expressed genes spanning pathways involved in cell-cycle regulation, endocrine signaling, immune function, oxidative stress, and lipid metabolism. Unexpectedly, more than half of the PFOA-induced genes were the same genes that normally increase during juvenile maturation, representing a highly significant enrichment of the endogenous developmental program. These findings indicate that the embryonic transcriptome is largely buffered against chronic PFOA exposure, whereas post-hatch stages exhibit heightened vulnerability. Rather than inducing a distinct toxicological state, PFOA predominantly acted by amplifying existing developmental gene regulatory programs. Our results identify the juvenile stage following hatching, equivalent to human neonatal and adolescent developmental stages, as a critical window of PFAS susceptibility and suggest that environmental contaminants may exert their effects by exaggerating normal developmental trajectories, with potential consequences for growth, maturation, and long-term health.

## Introduction

Per- and polyfluoroalkyl substances (PFAS) are a class of persistent synthetic chemicals that have become ubiquitous environmental contaminants, with detectable levels present in drinking water, food sources, wildlife, and human populations worldwide.^[1–3]^ Among them, perfluorooctanoic acid (PFOA) is one of the most extensively studied compounds due to its environmental persistence, bioaccumulation potential, and association with adverse health outcomes. PFOA has been detected in contaminated drinking water supplies and aquatic wildlife ^[4–9]^ at concentrations far exceeding the U.S. Environmental Protection Agency’s 2024 Maximum Contaminant Level (MCL) of 4 ng/L^[10, 11]^; for example, the Hoosick Falls, New York water supply averaged 534 ng/L during peak contamination.^[12]^ Human exposure begins early in life, as PFAS can readily cross the placenta and can also be transferred through breast/formula milk, resulting in detectable concentrations in fetuses, infants, and children.^[13–17]^

Growing epidemiological evidence suggests that early-life PFAS exposure may have lasting developmental consequences. Prenatal and neonatal PFAS exposure has been associated with developmental delays, including reduced language development, impaired motor function, altered neurobehavioral outcomes, and decreased cognitive performance during childhood.^[18–24]^ Several studies have further reported associations between PFAS exposure and altered growth trajectories, endocrine disruption, immune dysfunction, and reproductive abnormalities.^[25–31]^ Importantly, adolescence may represent an additional period of vulnerability. Recent epidemiological studies have linked PFAS exposure in teenagers and young adults with reduced lean muscle mass and bone density, altered body composition, and metabolic dysfunction, suggesting that PFAS may interfere with biological programs regulating growth, tissue maturation, and homeostasis during critical developmental transitions.^[13, 32, 33]^

Despite increasing evidence linking PFAS exposure to developmental abnormalities, a fundamental mechanistic question remains unresolved. Does PFOA impose a novel transcriptional program on developing organisms, or does it perturb endogenous developmental programs that are already underway? Distinguishing between these possibilities is critical for understanding how environmental contaminants influence developmental trajectories and long-term health outcomes.

The African turquoise killifish, *Nothobranchius furzeri*, provides a uniquely powerful vertebrate model for addressing this question across multiple developmental windows. As the shortest-lived captive vertebrate, with a median lifespan of approximately 3-7 months^[34]^, the killifish enables the study of developmental and aging processes within experimentally tractable timescales.^[35–37]^ Proportional lifespan scaling places the 1-day post-hatch juvenile at approximately the human infant stage and the 4-week post-hatch juvenile at adolescence^[38]^. The species reaches sexual maturity within 4-6 weeks of hatching^[39–41]^ and recapitulates many conserved vertebrate hallmarks of aging, including telomere shortening, neurodegeneration, chronic inflammation, and age-associated tumor formation^[42–44]^. Together with its annotated genome and growing use in environmental toxicology studies^[45]^, these features position *N. furzeri* as a valuable model for examining how environmental exposures interact with developmental and lifespan programs.

Here, we performed whole-transcriptome profiling of *N. furzeri* continuously exposed to an environmentally relevant concentration of PFOA (700 ng/L) from egg laying through juvenile development. By examining both newly hatched and older juvenile fish, we identify a striking stage-dependent response to chronic exposure. We demonstrate that embryonic developmental programs are largely buffered against PFOA-induced perturbation, whereas post-hatch juveniles exhibit extensive transcriptional remodeling across pathways regulating cell proliferation, endocrine signaling, immunity, oxidative stress, and metabolism. Unexpectedly, the majority of PFOA-responsive genes correspond to genes that normally change during juvenile maturation, indicating that PFOA predominantly acts by amplifying endogenous developmental gene regulatory programs rather than inducing a distinct toxicological state. These findings reveal a previously unrecognized mechanism of PFAS developmental toxicity and identify juvenile maturation as a critical window of susceptibility to environmental exposure.

## Materials and Methods

### Animal care and husbandry

African turquoise killifish, *Nothobranchius furzeri*, ZMZ strain, were maintained in a recirculating aquatic system (Aquaneering, San Diego, CA, USA) at 26–27°C under a 12 h light:12 h dark photoperiod. Water was prepared using reverse-osmosis (RO) water supplemented with Instant Ocean® sea salt and maintained at pH 7.0–7.5. Adult fish were fed live brine shrimp (Artemia, INVE Aquaculture Inc.) and supplemented with Otohime C1 (Reed Mariculture) frozen bloodworms (Hikari Bio-Pure Blood Worm Cubes). All animal procedures were approved by the Institutional Animal Care and Use Committee (IACUC) at North Carolina Central University (Protocol DK10112024).

Embryos were collected and maintained according to established killifish husbandry protocols with minor modifications.^[46]^ Embryo medium consisted of 1 L RO water supplemented with two Ringer’s tablets (MilliporeSigma, Cat. No. 96724), sterilized by autoclaving (121°C, 15 psi, 20 min), and supplemented with 0.002% methylene blue. Embryos were maintained in covered Petri dishes at 26°C until hatching.

### PFOA chemical exposure

Perfluorooctanoic acid (PFOA; Sigma-Aldrich, Cat. No. 171468-5G, ≥95% purity) was dissolved directly into embryo medium to achieve a final concentration of 700 ng/L. Embryos were continuously exposed from egg laying until tissue collection and maintained in 100 mm Petri dishes at 26°C. Control untreated embryos were maintained in embryo medium without PFOA. The exposure concentration was selected because it falls within the range reported in PFAS-contaminated communities and levels present in contaminated teleost fishes even though it exceeds the current U.S. EPA Maximum Contaminant Level (MCL) of 4 ng/L^[4–11]^. Embryo media without or without exposure to PFOA was replaced daily for all conditions.

### Embryo hatching and juvenile husbandry

Embryos displaying fully developed golden eyes were induced to hatch by transfer to 1.5-mL microcentrifuge tubes containing a minimal volume of embryo medium, with caps closed, and tubes kept at a 45° angle for 20 mins. Newly hatched juveniles were transferred to hatching solution overnight and subsequently moved to 0.8-L tanks containing a 1:1 mixture of embryo medium and RO water. Fish were fed live brine shrimp daily and maintained under a 12 h light:12 h dark cycle. After one week, juveniles were transitioned to RO water and housed in aerated 0.8-L tanks. Fast- and slow-growing juveniles were separated to minimize density-dependent growth effects. Water was changed daily in both control and PFOA-treated groups.

### RNA isolation and sequencing

To examine transcriptomic responses across development, fish were collected at two developmental stages: (1) 1-day post-hatching juveniles, corresponding to approximately four weeks of embryonic exposure, and (2) older juveniles collected after an additional four weeks of growth, corresponding to approximately eight weeks of total exposure. For each developmental stage and treatment group, three biological replicates consisting of an individual juvenile fish were flash frozen and processed independently.

Whole animals were homogenized in 50 μL TRIzol Reagent (Invitrogen, Cat. No. 15596018), and total RNA was isolated using the Direct-zol RNA Miniprep Kit (Zymo Research, Cat. No. R2073) according to the manufacturer’s instructions, including on-column DNase treatment. RNA concentration and purity were assessed using a NanoDrop One spectrophotometer (Thermo Fisher Scientific), and all samples exhibited A260/A280 ratios greater than 1.8.

RNA samples were submitted to Novogene (Sacramento, CA, USA) for library preparation and sequencing. Libraries were generated using the Illumina TruSeq Stranded mRNA Library Preparation Kit and sequenced on the Illumina NovaSeq platform to produce 150-bp paired-end reads. Sequencing yielded 270–403 Gb per sample with Q30 values ranging from 79–85%, indicating high-quality sequence data.

### Bioinformatics and statistical analysis

Raw sequencing reads were aligned to the *N. furzeri* reference genome (GCF_001465895.1_Nfu_20140520) using HISAT2. Gene-level read counts were generated and imported into R for downstream analysis. Differential gene expression analysis was performed using the edgeR package. Lowly expressed genes were filtered prior to analysis, and library sizes were normalized using the trimmed mean of M-values (TMM) method. Gene-wise dispersions were estimated using a negative binomial model, and differential expression was assessed using generalized linear models implemented in edgeR. P-values were corrected for multiple hypothesis testing using the Benjamini–Hochberg false discovery rate (FDR) procedure. Genes with an adjusted p-value (FDR) ≤ 0.05 and an absolute log2 fold change ≥ 2.0 were considered significantly differentially expressed. A total of 24,495 annotated mRNA features were evaluated. Four planned comparisons (C) were performed: (C1) PFOA-exposed versus untreated 1-day hatched juveniles, (C2) PFOA-exposed versus untreated older juveniles, (C3) untreated older juveniles versus untreated 1-day hatched juveniles to define the endogenous developmental maturation program, and (C4) PFOA-exposed older juveniles versus PFOA-exposed 1-day hatched juveniles. Within-group Pearson R^2^ correlations were: untreated 1-day hatched juvenile 0.979 (range 0.967–0.989); PFOA exposed 1-day hatched juvenile 0.988 (range 0.985–0.992); untreated older juvenile 0.982 (range 0.979–0.987); PFOA exposed older juvenile 0.977 (range 0.970–0.989), confirming high biological reproducibility. A total of 24,495 mRNA features were assessed genome-wide across four planned comparisons. Gene Ontology (GO) and Kyoto Encyclopedia of Genes and Genomes (KEGG) enrichment analyses were conducted using the clusterProfiler package, with significance defined as FDR-adjusted p ≤ 0.05. All statistical analyses and data visualizations were performed in R.

### Hypergeometric enrichment analysis

To determine whether PFOA-responsive genes preferentially overlapped with genes associated with normal developmental maturation, a hypergeometric enrichment test was performed. Genes significantly upregulated during normal maturation were identified from comparison C3 (untreated older juveniles versus untreated 1-day hatched juveniles). The overlap between these genes and genes upregulated following PFOA exposure in older juveniles (C2) was evaluated relative to the total number of genes tested. The hypergeometric distribution was parameterized as follows: total gene population (M) = 24,495, developmental maturation genes (K) = 485, PFOA-upregulated genes (n) = 427, and observed overlap (k) = 249. Statistical significance was calculated in R using the function “phyper(k−1, K, M−K, n, lower.tail = FALSE).” An analogous analysis was performed for PFOA-downregulated genes. Plots were made in R using ggplot or obtained from NovoMagic: Online RNA-seq Bioinformatics Analysis Tool from Novogene.

## Results

### PFOA exposure elicits a stage-dependent transcriptomic response during African killifish, *Nothobranchius furzeri*, development

To determine how susceptibility to PFOA exposure changes across development, African killifish, *Nothobranchius furzeri*, were continuously exposed to 700 ng/L PFOA from egg laying through juvenile development and sampled at two developmental stages: 1-day post-hatching juveniles, corresponding approximately to the human infant stage, and older juveniles, corresponding to adolescence (Figure 1A). Whole-transcriptome RNA sequencing was performed to characterize molecular responses to chronic exposure.

**Figure 1.**
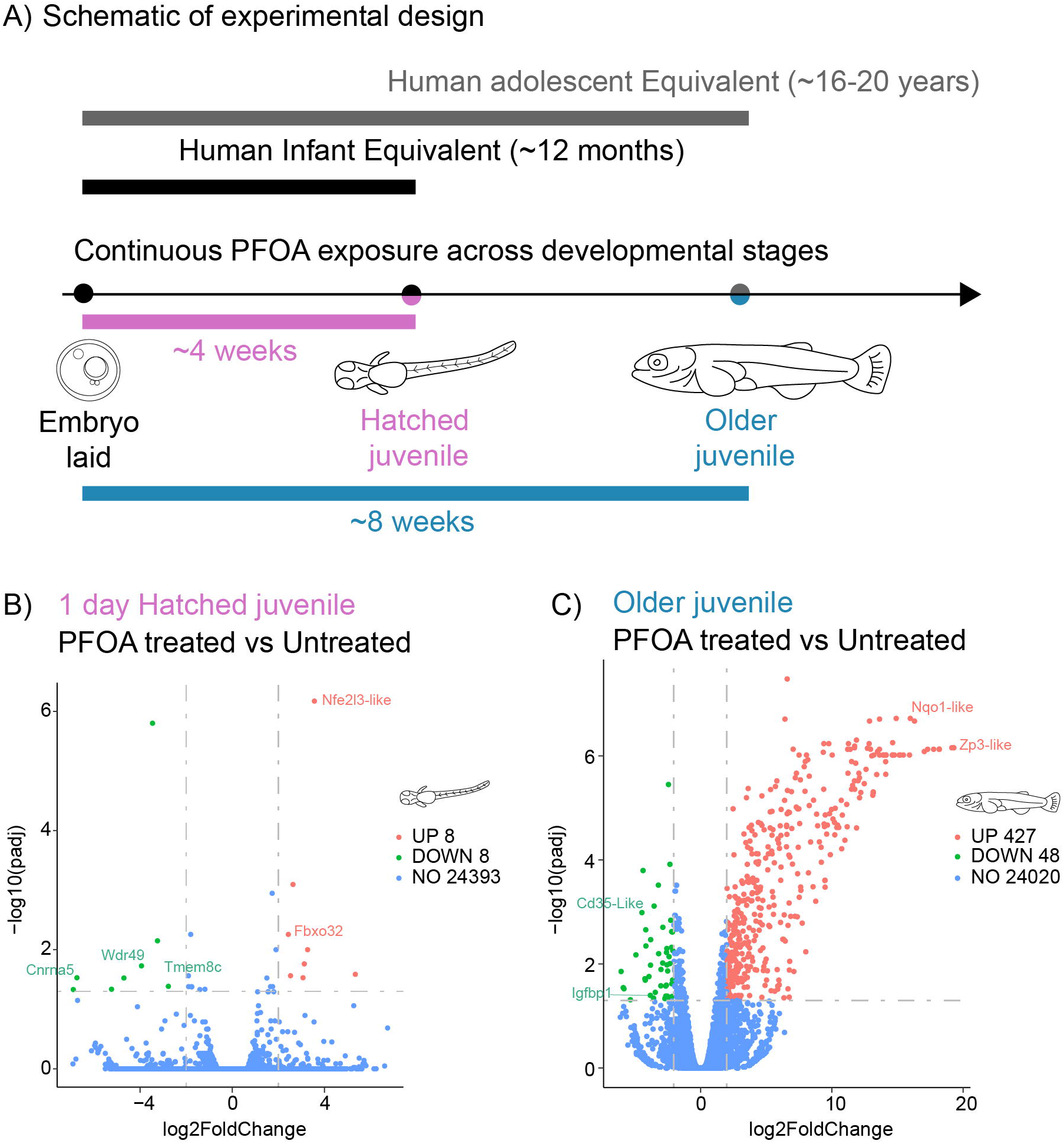
Stage-dependent transcriptomic responses to continuous PFOA exposure during killifish development. A) Experimental design. African killifish (*Nothobranchius furzeri*) embryos were continuously exposed to 700 ng/L PFOA from egg laying through juvenile development. Samples were collected at two developmental stages: 1-day hatched juveniles (~4 weeks of exposure), corresponding approximately to the human infant stage, and older juveniles (~8 weeks of exposure), corresponding approximately to adolescence. B) Volcano plot showing differentially expressed genes (DEGs) between PFOA-exposed and untreated 1-day hatched juveniles. Significantly upregulated genes are shown in red, downregulated genes in green, and non-significant genes in blue. Differential expression was defined as adjusted p ≤ 0.05 and |log_2_FC| ≥ 2.0. C) Volcano plot showing DEGs between PFOA-exposed and untreated older juveniles following approximately 8 weeks of continuous exposure.

Despite approximately four weeks of continuous PFOA exposure throughout embryogenesis, 1-day hatched juveniles exhibited a remarkably restricted transcriptional response when compared with untreated controls (Figure 1B). Differential expression analysis identified only 16 significantly altered mRNAs, consisting of eight upregulated and eight downregulated genes. This limited response was unlikely to reflect insufficient statistical power, as biological replicates exhibited high reproducibility (Pearson R^2^ = 0.985–0.992), the same samples yielded 50 differentially expressed miRNAs in parallel small RNA sequencing analyses, and several transcripts displayed highly significant expression changes.

The most significantly upregulated gene was *nfe2l3-like* (log_2_FC = +3.56, padj = 6.7 × 10^−7^), a transcription factor associated with integrated stress responses. Additional upregulated genes included *arg2* (arginase 2; log_2_FC = +2.53) and *fbxo32* (F-box protein 32/atrogin-1; log_2_FC = +2.43), whereas the most strongly downregulated transcript was *chrna5* (nicotinic acetylcholine receptor alpha-5; log_2_FC = −6.74). Consistent with the limited number of transcriptional changes, KEGG pathway analysis identified few significantly enriched pathways (Supp Figure 1), including FoxO signaling pathway and arginine biosynthesis and arginine-proline metabolism (padj < 0.05). Together, these findings suggest that the embryonic developmental program is largely buffered against broad transcriptomic disruption by chronic PFOA exposure, with activation of stress signaling representing the primary molecular response.

In contrast, transcriptomic responses expanded dramatically in older juveniles exposed to PFOA for approximately eight weeks (Figure 1C). Differential expression analysis identified 475 significantly altered genes, including 427 upregulated and 48 downregulated transcripts, representing an approximately 30-fold increase in the number of differentially expressed genes relative to newly hatched juveniles. The predominance of upregulated transcripts suggests that chronic PFOA exposure primarily activates gene expression programs rather than broadly suppressing transcription. Notably, oxidative stress-associated genes, including *nqo1-like*, were among the most highly induced transcripts, indicating persistent activation of cellular stress pathways during juvenile development. In contrast, *igfbp1*, a gene involved in insulin-like growth factor signaling and the positive regulation of cell growth, was among the downregulated transcripts, indicating potential disruptions in growth and metabolic regulation.

The striking difference in transcriptional responses between developmental stages demonstrates that susceptibility to PFOA exposure is strongly stage dependent. While embryos and newly hatched juveniles appear largely resistant to widespread transcriptomic perturbation, older juveniles exhibit extensive transcriptional remodeling. These findings identify post-hatching juvenile development as a critical window of PFAS susceptibility and suggest that the biological consequences of chronic exposure emerge progressively during maturation rather than during embryogenesis alone.

### Chronic PFOA exposure remodels developmental, metabolic, and stress-response pathways in older juveniles

Given the pronounced increase in transcriptional responsiveness observed in older juveniles, we next examined the biological processes and pathways disrupted by 8 weeks of chronic PFOA exposure. Hierarchical clustering of the 475 differentially expressed genes revealed clear separation between untreated and PFOA-exposed animals, representing a 30-fold increase relative to the 1-day hatched juvenile stage, and indicating a robust and coordinated transcriptional response to exposure (Figure 2A).

**Figure 2.**
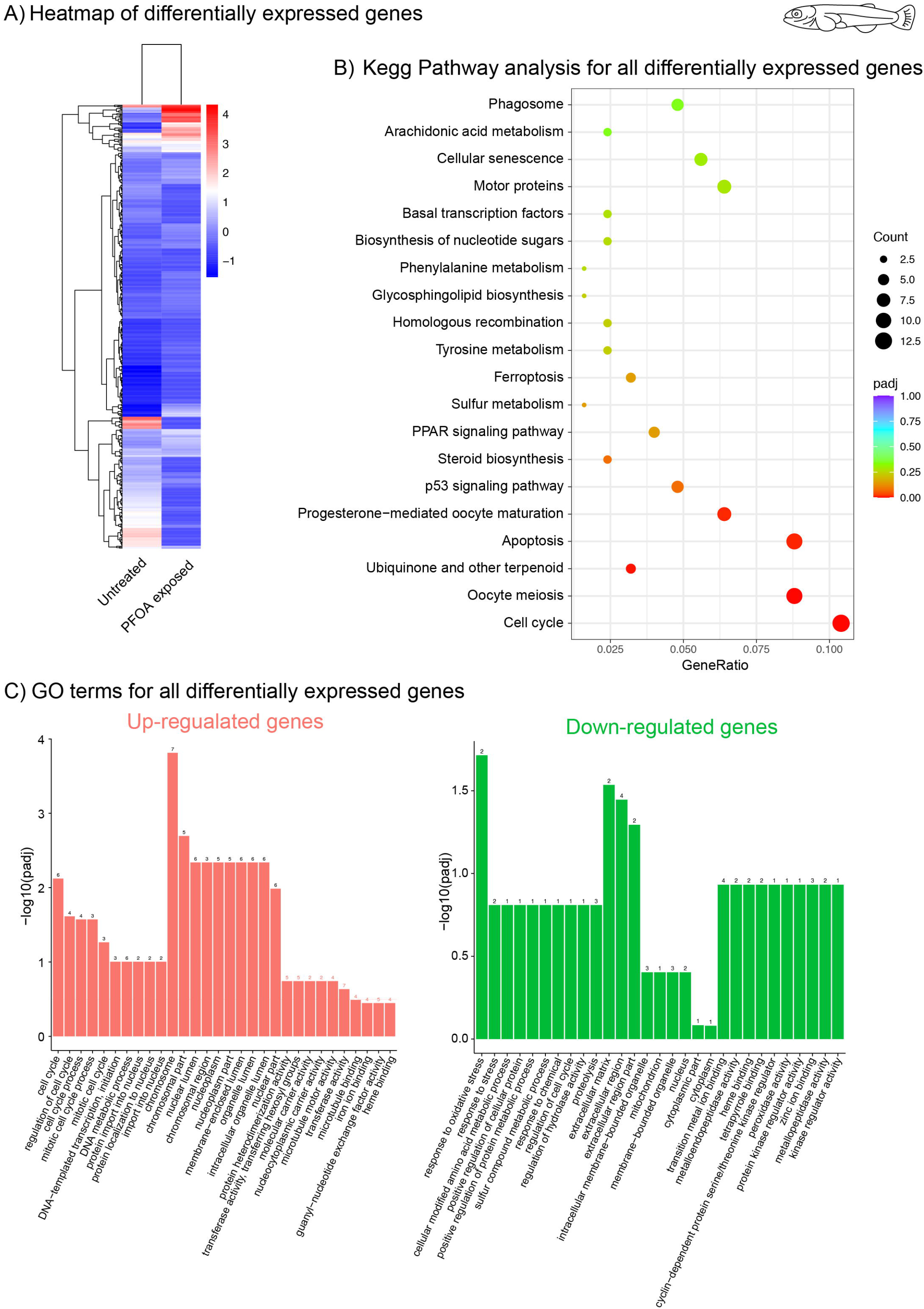
Chronic PFOA exposure remodels developmental, metabolic, and stress-response pathways in older juveniles. A) Hierarchical clustering heatmap of the 475 differentially expressed genes identified between untreated and PFOA-exposed older juveniles. Gene expression values are represented as scaled expression (Z-score). B) KEGG pathway enrichment analysis of differentially expressed genes in older juveniles exposed to PFOA. Dot size represents the number of genes associated with each pathway, and color indicates adjusted p-value. C) Gene Ontology (GO) enrichment analysis of upregulated genes (left) and downregulated genes (right). Bar height indicates –log10(adjusted p-value).

KEGG pathway enrichment analysis identified multiple significantly enriched pathways associated with cellular proliferation, metabolism, stress responses, and developmental regulation (Figure 2B). Among the most significantly enriched pathways were cell cycle, oocyte meiosis, progesterone-mediated oocyte maturation, and p53 signaling, suggesting widespread alterations in mechanisms governing cell proliferation, growth, and developmental progression. Additional enrichment of apoptosis, cellular senescence, and ferroptosis pathways indicates activation of cellular stress and damage-response mechanisms. Metabolic pathways affected by exposure included steroid biosynthesis, PPAR signaling, arachidonic acid metabolism, tyrosine metabolism, phenylalanine metabolism, and sulfur metabolism, highlighting broad disruption of lipid, hormone, and energy homeostasis. Enrichment of phagosome and homologous recombination pathways further suggests alterations in immune function and genome maintenance.

Gene Ontology (GO) analysis of upregulated genes revealed significant enrichment for biological processes associated with cell proliferation and developmental progression (Figure 2C, left panel). Enriched terms included cell cycle, regulation of the cell cycle, mitotic cell-cycle processes, DNA-templated transcription initiation, DNA replication, protein localization to the nucleus, chromosome organization and segregation, and developmental growth. Upregulated genes were also associated with ribosomal and chromosomal components, membrane-enclosed lumen organization, and intracellular organelle functions, indicating increased transcriptional, biosynthetic, and proliferative activity in response to chronic PFOA exposure.

In contrast, downregulated genes were enriched for processes involved in oxidative stress responses, amino acid metabolism, sulfur compound metabolism, regulation of proteolysis, and extracellular matrix-related functions (Figure 2C, right panel). Additional enrichment was observed for cellular and extracellular structures, including extracellular matrix components, membrane-bounded organelles, and cytoplasmic structures. Molecular function terms associated with metallopeptidase activity, tetrapyrrole binding, peroxidase activity, and kinase regulatory functions were also reduced, suggesting suppression of pathways involved in stress adaptation, metabolism, and cellular signaling.

Together, these analyses indicate that chronic PFOA exposure induces broad transcriptional remodeling in juvenile fish. The simultaneous upregulation of genes involved in cell-cycle progression, growth, and chromosomal organization, coupled with downregulation of oxidative stress response, metabolic, and extracellular matrix-related pathways, suggests a shift toward proliferative programs at the expense of normal cellular homeostasis and stress-response mechanisms.

### PFOA amplifies endogenous developmental programs across multiple biological pathways

To determine whether PFOA-induced transcriptional responses reflected activation of novel toxicological pathways or amplification of normal developmental programs, we compared genes altered during juvenile maturation in untreated fish with those changing during maturation under continuous PFOA exposure. Differential expression analysis identified 588 maturation-associated genes in untreated fish, consisting of 103 upregulated and 485 downregulated transcripts (Figure 3A). In contrast, maturation under continuous PFOA exposure was associated with substantially greater transcriptional remodeling, with 1,021 differentially expressed genes, including 458 upregulated and 563 downregulated transcripts (Figure 3B).

**Figure 3.**
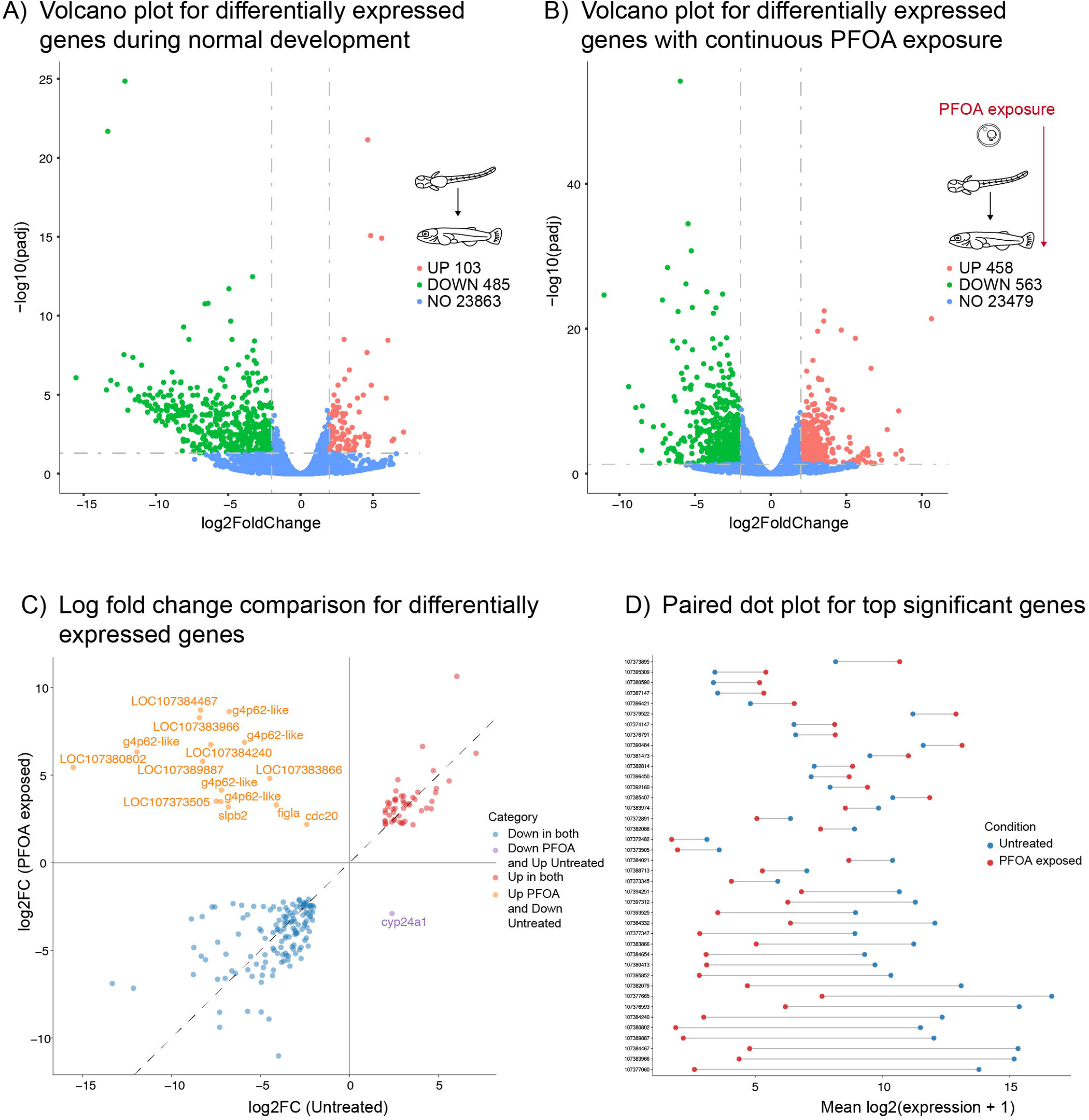
PFOA amplifies endogenous developmental maturation programs. A) Volcano plot showing genes differentially expressed during normal juvenile maturation, comparing untreated older juveniles with untreated 1-day hatched juveniles. B) Volcano plot showing genes differentially expressed during maturation under continuous PFOA exposure, comparing PFOA-exposed older juveniles with PFOA-exposed 1-day hatched juveniles. C) Comparison of log_2_ fold-change values for genes differentially expressed during normal maturation and maturation under PFOA exposure. Genes exhibiting concordant regulation between developmental trajectories are shown along the diagonal, indicating preservation of developmental directionality under exposure. D) Paired expression plot for representative highly significant genes demonstrating differences in expression between untreated and PFOA-exposed developmental trajectories.

Direct comparison of developmental trajectories revealed a strong positive relationship between gene expression changes occurring during normal maturation and those occurring during maturation under PFOA exposure (Figure 3C). The majority of genes exhibited concordant directions of regulation between datasets, indicating that developmental progression remains largely intact despite chronic exposure. Hierarchical clustering analyses similarly demonstrated clear separation between hatched and older juveniles in both untreated and PFOA-exposed animals (Supp. Figure 2 A,B), while GO term enrichment analyses revealed substantial overlap in the biological processes associated with maturation under both conditions (Supp. Figure 2 C,D). Together, these findings suggest that PFOA does not fundamentally redirect developmental trajectories but instead enhances existing developmental gene regulatory programs.

Comparison of shared maturation-associated genes further demonstrated that most overlapping transcripts were regulated in the same direction in both untreated and PFOA exposed animals (Supp. Figure 3A). Genes involved in extracellular and secreted protein functions, including lectin family members such as L-rhamnose-binding lectin CSL2-like, nattectin-like lectins, and sialic acid-binding lectins, were consistently upregulated in both datasets, suggesting common changes in extracellular signaling and host-defense pathways during juvenile maturation. In contrast, genes associated with proteolysis and protein turnover, including proteasome subunits (e.g., psmb8), ubiquitin pathway regulators (trim21, trim39, arih2, and mib2), and serine proteases such as elastase-1-like genes, were consistently downregulated, indicating reduced protein degradation and proteostatic activity with age. Notably, while these shared patterns likely reflect normal developmental maturation, PFOA exposure was additionally associated with altered regulation of oxidative stress and metabolic genes, suggesting that environmental exposure modifies age-related transcriptional programs by disrupting pathways involved in cellular stress responses and metabolic homeostasis. Furthermore, a small subset of genes, including cyp24a1, exhibited attenuated or opposing expression patterns between the two datasets, indicating that PFOA selectively alters specific developmental and metabolic pathways during juvenile maturation.

To formally quantify the relationship between PFOA-responsive genes and endogenous developmental programs, we performed a hypergeometric enrichment analysis. Of the 427 genes upregulated by PFOA in older juveniles, 249 genes (58.3%) were also among the genes normally upregulated during maturation. This overlap was highly significant, corresponding to a 29.5-fold enrichment over random expectation (hypergeometric test, p < 2.2 × 10^−366^). Enrichment among downregulated genes was substantially weaker, indicating that the dominant effect of PFOA is enhancement of developmental gene activation rather than repression.

Together, these findings demonstrate that chronic PFOA exposure does not induce a distinct toxicological transcriptional program. Instead, PFOA predominantly acts by amplifying endogenous developmental gene regulatory programs that normally accompany juvenile maturation, resulting in exaggerated developmental trajectories that converge on interconnected biological axes of cell-cycle regulation, endocrine signaling, immune function, oxidative stress responses, and lipid metabolism.

## Discussion

PFAS exposure has been associated with developmental, neurological, metabolic, and immune-related health outcomes, yet the mechanisms by which these contaminants alter developmental trajectories remain poorly understood. Using the African killifish (*Nothobranchius furzeri*), we demonstrate that susceptibility to chronic PFOA exposure is highly stage dependent. While embryos and newly hatched juveniles exhibited remarkably limited transcriptional responses despite four weeks of continuous exposure, older juveniles displayed extensive transcriptomic remodeling after eight weeks of exposure. These findings identify post-hatching juvenile development as a critical window of PFAS susceptibility.

A major finding of this study is that PFOA does not primarily induce a novel toxicological transcriptional program. Instead, more than half of the genes upregulated by PFOA were also genes that normally increase during juvenile maturation, representing a highly significant enrichment of the endogenous developmental program. Furthermore, most shared genes changed in the same direction in both untreated and exposed animals, indicating that developmental progression remains intact but is exaggerated under exposure. Together, these results suggest that PFOA acts as an amplifier of existing developmental gene regulatory programs rather than redirecting development toward an alternative transcriptional state.

This amplification was particularly evident in the zona pellucida gene cluster, where genes that normally increase during maturation exhibited substantially greater induction under PFOA exposure. Similar patterns were observed among genes involved in cell-cycle progression and developmental growth, suggesting that PFOA disrupts mechanisms that normally constrain developmental gene expression. Such a mechanism provides a new perspective on developmental toxicity, whereby environmental contaminants alter the magnitude of developmental programs rather than simply activating stress-response pathways.

Pathway analyses further revealed that amplified developmental trajectories converge on major biological axes of cell-cycle regulation, endocrine signaling, immune function, oxidative stress responses, and lipid metabolism. These findings are consistent with epidemiological and experimental studies linking PFAS exposure to altered growth, endocrine disruption, immune dysfunction, and metabolic abnormalities. Notably, oxidative stress responses progressed from modest activation of *nfe2l3-like* in newly hatched juveniles to robust induction of *nqo1-like* genes in older juveniles, suggesting a transition from adaptive stress sensing to sustained cellular stress during prolonged exposure.

Our findings also have implications for understanding developmental windows of vulnerability. Although embryonic development is often considered the most sensitive period for environmental exposures, the relatively limited embryonic response observed here suggests that developmental buffering mechanisms may protect early embryos from widespread transcriptional disruption. In contrast, juvenile maturation appears substantially more susceptible, consistent with epidemiological studies linking PFAS exposure during childhood and adolescence to altered growth, neurodevelopment, and reduced lean muscle mass.

In summary (Figure 4), chronic PFOA exposure produces highly stage-dependent effects during vertebrate development. While embryonic developmental programs appear largely buffered against transcriptional disruption, juvenile fish exhibit widespread remodeling of gene expression. Rather than inducing an entirely distinct toxicological state, PFOA primarily amplifies endogenous developmental gene regulatory programs, resulting in exaggerated maturation-associated trajectories across multiple biological systems. We propose a model in which early embryonic exposure elicits a modest oxidative stress response while intrinsic developmental buffering mechanisms maintain transcriptional stability. As animals transition into juvenile stages under continuous PFOA exposure, however, these protective mechanisms become insufficient, leading to enhanced activation of pathways involved in cell-cycle regulation, endocrine signaling, immune function, oxidative stress responses, and lipid metabolism.

**Figure 4.**
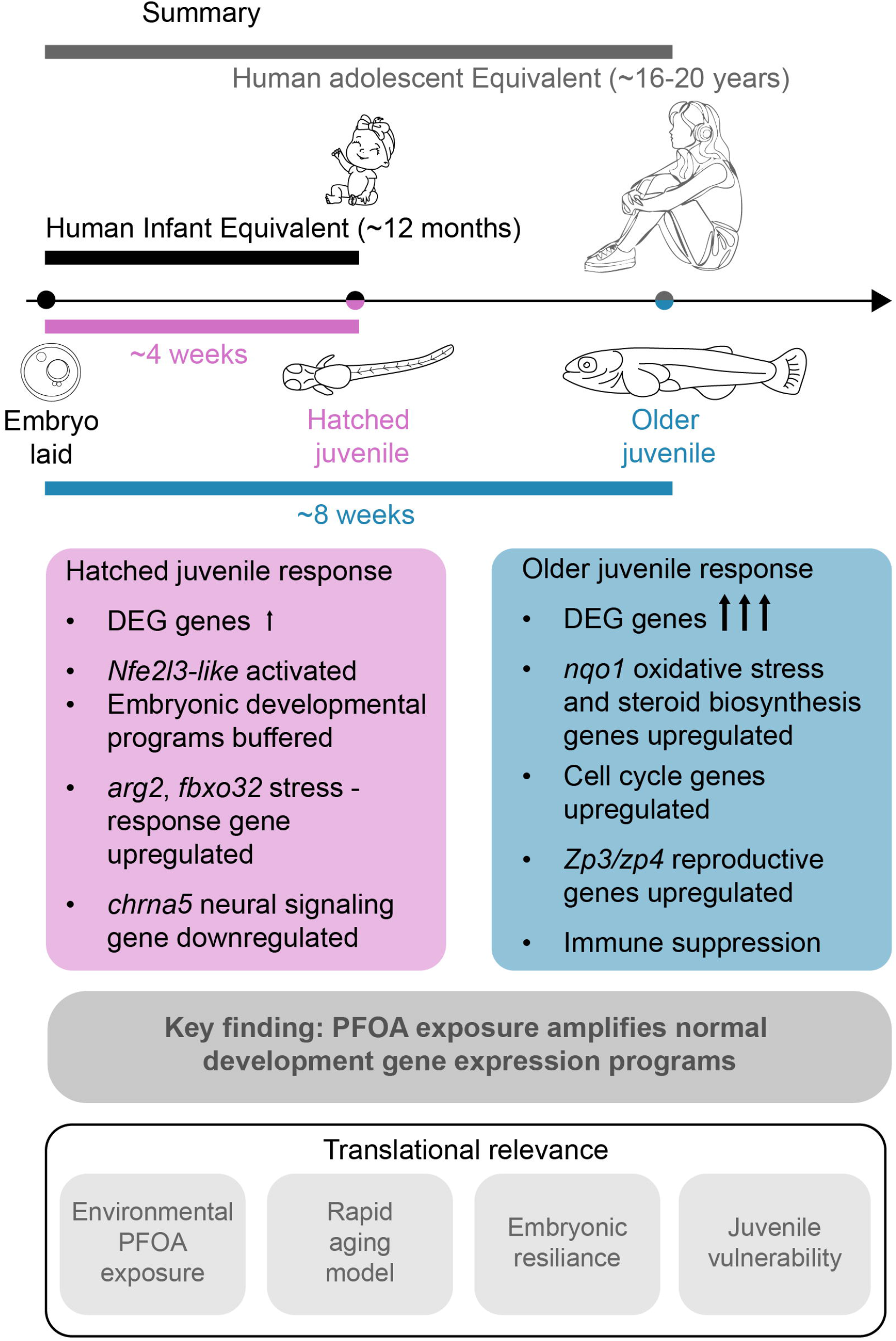
Proposed model for stage-dependent PFOA susceptibility and amplification of developmental gene expression programs. Continuous exposure to an environmentally relevant concentration of PFOA (700 ng/L) produces distinct transcriptional responses across developmental stages. Newly hatched juveniles (~4 weeks of exposure) exhibit limited transcriptomic disruption characterized by activation of stress-response genes. In contrast, older juveniles (~8 weeks of exposure) display extensive transcriptional remodeling. Rather than inducing a novel toxicological program, PFOA predominantly amplifies endogenous developmental maturation pathways.

Importantly, the exposure concentration used in this study (700 ng/L PFOA) is within the range reported in contaminated drinking water sources, increasing the environmental and translational relevance of these findings. The heightened sensitivity observed during the post-hatching juvenile period suggests that susceptibility to PFAS exposure extends beyond embryogenesis and may represent a critical window of vulnerability. Given that this developmental stage broadly parallels childhood and adolescence in humans, our findings support the need to also consider postnatal developmental periods in PFAS risk assessment. Collectively, these results provide a new framework for understanding PFAS developmental toxicity, suggesting that environmental contaminants may exert long-term effects not only by disrupting development, but also by amplifying normal developmental programs during critical windows of maturation.

## Acknowledgements

This research was supported by the RCMI Program grant U54MD012392 awarded to DK, an RCMI Pilot Grant awarded to ZA, and funding from the NC PFAS Collaboratory through the project Molecular Mechanisms of PFAS. We acknowledge the Julius L. Chambers Biomedical/Biotechnology Research Institute (BBRI) at North Carolina Central University for providing research facilities and resources. We thank the BBRI faculty and staff for their support, particularly Dr. Derek Norford, Ms. Camilla Felton, Thomas Vanden Broek, Dr. Qing Cheng, and Dr. Claudia Alberico. We are grateful to Dr. Anne Brunet and members of the Brunet Laboratory at Stanford University for their guidance in establishing the killifish facility and colony. We also thank Riley Galton and Robert Schnittker from the Sánchez Alvarado Laboratory at the Stowers Institute for all their support and expertise in guiding us for killifish embryo and juvenile husbandry, and Wouter Lanneau of Microbiotests for his assistance with killifish embryos. We also acknowledge the use of an AI agent for discussion and interpretation of the study findings.

## Figure legends

**Supp Figure 1** Transcriptomic response to PFOA exposure in 1-day post-hatch juveniles. A) Hierarchical clustering heatmap of genes differentially expressed between PFOA-exposed and untreated 1-day post-hatch juveniles. Gene expression values are represented as scaled expression (Z-score). B) KEGG pathway enrichment analysis of differentially expressed genes between PFOA-exposed and untreated 1-day post-hatch juveniles. C) Gene Ontology (GO) enrichment analysis of differentially expressed genes between PFOA-exposed and untreated 1-day post-hatch juveniles.

**Supp Figure 2** Developmental maturation programs in untreated and PFOA-exposed juveniles. A) Hierarchical clustering heatmap of genes differentially expressed between untreated 1-day post-hatch juveniles and untreated older juveniles. B) Hierarchical clustering heatmap of genes differentially expressed between PFOA-exposed 1-day post-hatch juveniles and PFOA-exposed older juveniles. C) Gene Ontology enrichment analysis of maturation-associated genes identified in untreated animals. D) Gene Ontology enrichment analysis of maturation-associated genes identified in PFOA-exposed animals.

**Supp Figure 3** Shared and discordant maturation-associated genes between untreated and PFOA-exposed developmental trajectories. A) Heatmap showing genes significantly altered during maturation in both untreated and PFOA-exposed animals. Color intensity represents log_2_ fold change values. B) Heatmap showing genes exhibiting discordant regulation between untreated and PFOA-exposed developmental trajectories.

## Notes

### Competing Interest Statement

The authors have declared no competing interest.

