## Supplementary figures and images for "PFOA exposure amplifies normal developmental gene expression programs in the African Killifish, *Nothobranchius furzeri*"

### Supp Figure 1

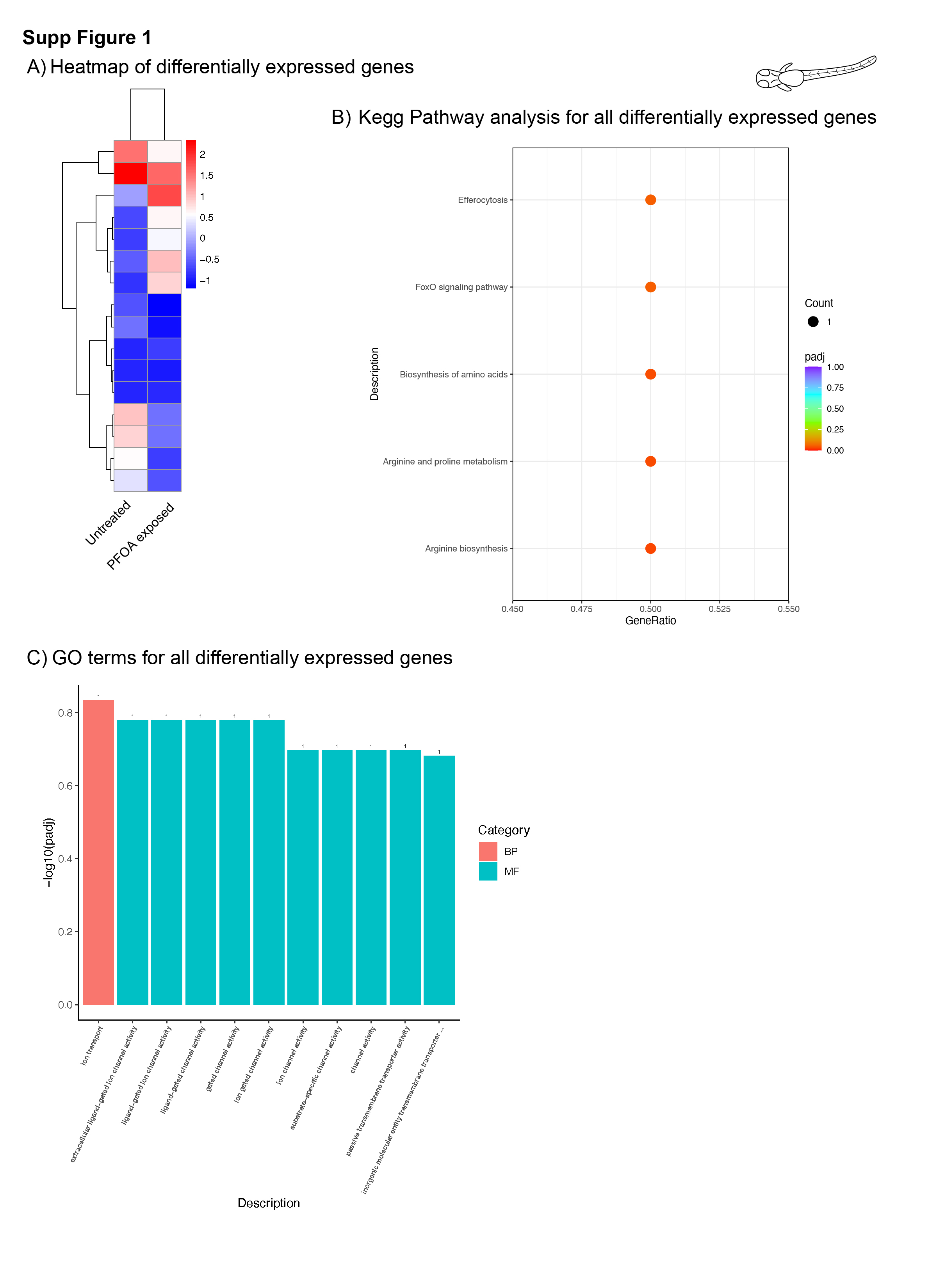

### Supp Figure 2

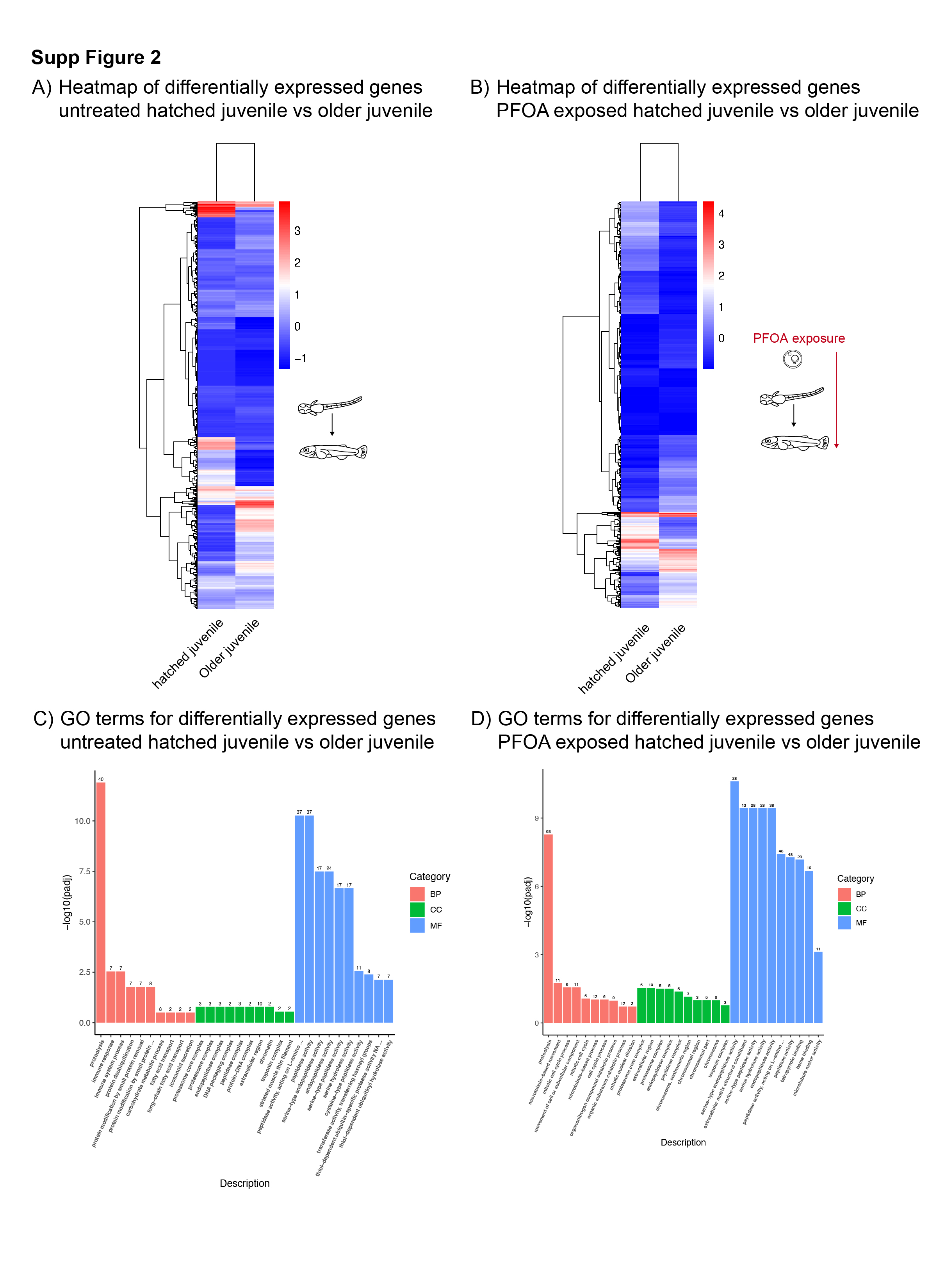

### Supp Figure 3

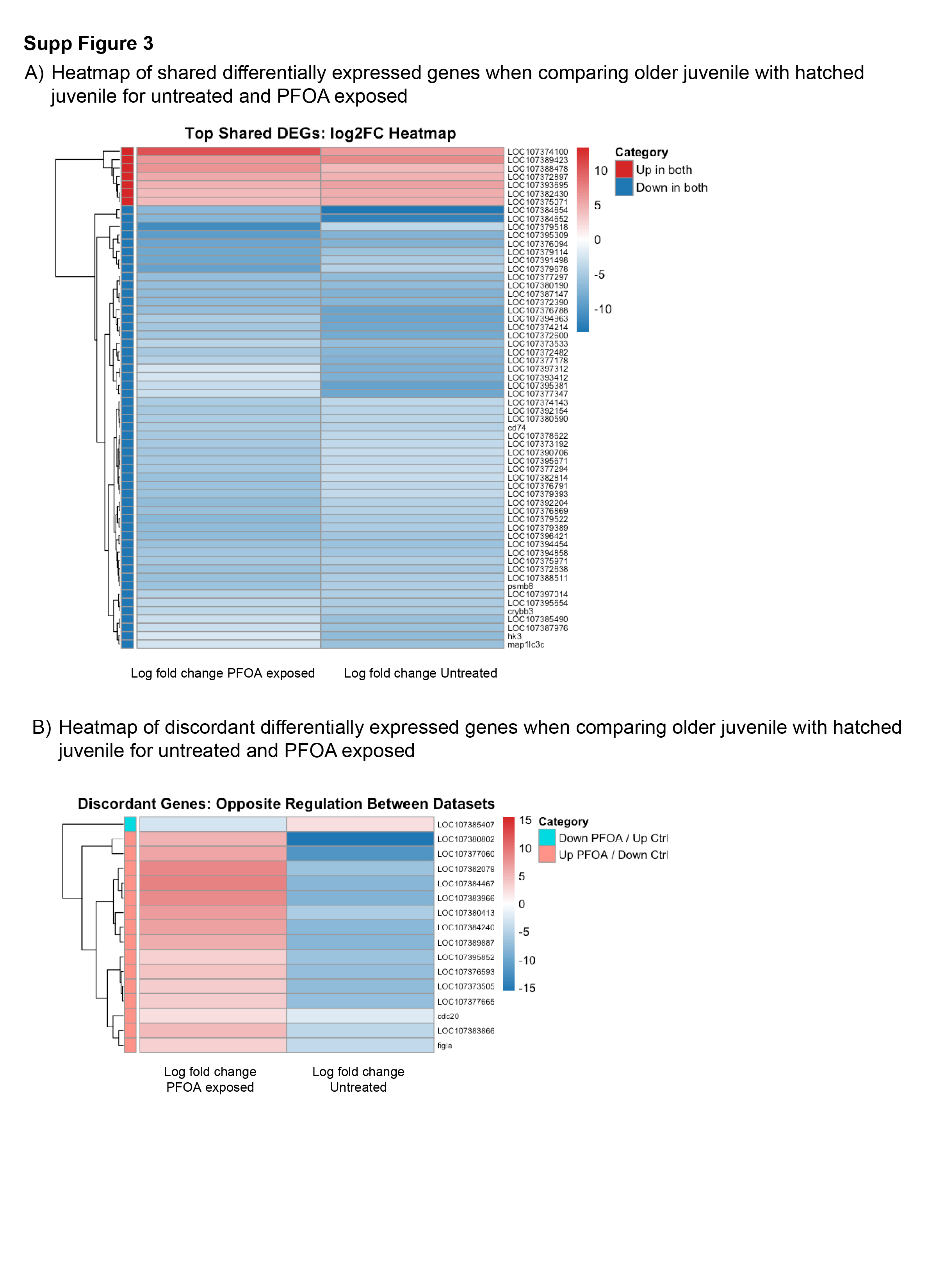
